# The Intestinal Microbiota Contributes to the Control of Highly Pathogenic H5N9 Influenza Virus Replication in Ducks

**DOI:** 10.1101/778258

**Authors:** Thomas Figueroa, Pierre Bessière, Amelia Coggon, Roosmarijn van der Woude, Maxence Delverdier, Monique H. Verheije, Robert P. de Vries, Romain Volmer

## Abstract

Ducks usually show little or no clinical signs following highly pathogenic avian influenza virus infection. In order to analyze if the gut microbiota could contribute to the control of influenza virus replication in ducks, we used a broad-spectrum oral antibiotic treatment to deplete the gut microbiota before infection with a highly pathogenic H5N9 avian influenza virus. Antibiotic-treated ducks and non-treated control ducks did not show any clinical signs following H5N9 virus infection. We did not detect any difference in virus titers neither in the respiratory tract, nor in the brain and spleen. However, we found that antibiotic-treated H5N9 virus infected ducks had significantly increased intestinal virus excretion at day 3 and 5 post-infection. This was associated with a significantly decreased antiviral immune response in the intestine of antibiotic-treated ducks. Our findings highlight the importance of an intact microbiota for an efficient control of avian influenza virus replication in ducks.

**IMPORTANCE:** Ducks are frequently infected with avian influenza viruses belonging to multiple subtypes. They represent an important reservoir species of avian influenza viruses, which can occasionally be transmitted to other bird species or mammals, including humans. Ducks thus have a central role in the epidemiology of influenza virus infection. Importantly, ducks usually show little or no clinical signs even following infection with a highly pathogenic avian influenza virus. We provide evidence that the intestinal microbiota contributes to the control of influenza virus replication in ducks by modulating the antiviral immune response. Ducks are able to control influenza virus replication more efficiently when they have an intact intestinal microbiota. Therefore, maintaining a healthy microbiota by limiting perturbations to its composition should contribute to prevention of avian influenza virus spread from the duck reservoir.

## INTRODUCTION

Ducks have a central role in the epidemiology of influenza virus (1, 2). They have been shown to shed viruses belonging to multiple subtypes and represent the main reservoir of influenza viruses that can occasionally be transmitted to other animal species, including gallinaceous poultry, and occasionally mammals. Low pathogenic avian influenza viruses (LPAIVs) of the H5 and H7 subtype have the capacity to evolve to highly pathogenic avian influenza viruses (HPAIV). This evolution is due to the acquisition of a polybasic cleavage site at the hemagglutinin (HA) cleavage site, allowing the HA to be cleaved by intracellular ubiquitous proteases and the virus to spread systematically (3). By contrast, the HA of LPAIV is proteolytically matured by trypsin-like proteases expressed in the respiratory and digestive tract, thus restricting replication of LPAIV to these tissues in birds.

In chickens, HPAIV infection can reach 100% mortality in a few days, whereas ducks usually only exhibit mild clinical signs, if any, following HPAIV infection (4). The mechanisms contributing to the host species-dependent differences in pathogenicity are not fully understood. Ducks express the viral RNA sensor retinoic acid-induced gene I (RIG-I), which is absent from chickens (5). Melanoma differentiation-associated protein 5 (MDA-5) could compensate for the lack of RIG-I in chickens cells and mediate type I IFN responses to influenza A virus infection (6). However, comparative studies have provided evidence that HPAIV infections are associated with a more rapid type I interferon immune response in ducks compared to chickens and reduced pro-inflammatory cytokines expression in ducks compared to chickens. This may lead to reduced immunopathology and therefore less clinical signs in ducks (7–11). Extensive tissue damage due to higher levels of viral replication and inflammation in multiple organs are probably the cause of the higher HPAIV-associated mortality in chickens compared to ducks.

The gut microbiota is now recognized as a regulator of many physiological functions in the host, reaching far beyond its contribution to digestion (12). Several studies have highlighted the role of the gut microbiota in shaping the antiviral immune response against influenza virus, in mammals, in which the virus replicates in the respiratory tract (13–17), as well as following LPAIV infection in chickens, in which the virus replicates in the respiratory and digestive tract (18, 19).

Altogether, these observations prompted us to investigate to what extent the gut microbiota could contribute to the control of HPAIV infection in ducks. By treating ducks with a broad-spectrum antibiotic (ABX) cocktail, we achieved a significant depletion of the intestinal microbiota. Groups of non-treated and ABX-treated ducks were infected with a H5N9 HPAIV. We revealed that ABX-treated ducks had significantly higher viral excretion in the intestine, which correlated with an impaired antiviral immune response. This study thus demonstrates that the intestinal microbiota contributes to the control of avian virus replication in ducks.

## RESULTS

### Antibiotic treatment leads to a significant depletion of the intestinal microbiota

From two weeks of age, ABX-treated ducks were continuously treated with a broad-spectrum ABX cocktail though their drinking water (Fig. 1A). To assess the impact of the ABX-treatment on the intestinal microbiota, we determined the bacterial load from fresh feces harvested from non-treated control ducks and from ABX-treated ducks. After two weeks of ABX-treatment, we detected a greater than 10^6^-fold depletion of the intestinal bacteria cultivable under aerobic conditions (Fig. 1B), as well as a greater than 10^2^-fold depletion of the number of bacterial 16S rRNA gene copies (Fig. 1C). Using 16S rRNA FISH, we detected bacteria aggregating as filaments in the vicinity of intestinal epithelial cells on ileal tissue section from non-treated control ducks (Fig. 1D). In contrast, no bacteria were detected by FISH in ileal samples originating from ABX-treated ducks. Based on the culture method and culture independent methods, we can thus conclude that the ABX treatment caused a significant reduction in the ducks’ gut microbiota. The ABX treatment did not induce any clinical signs or change in the ducks’ behavior, nor did it diminish the quantity of water consumed. There was no significant difference in the average weight of the ABX-treated and the non-treated animals (data not shown).

**Figure 1.**
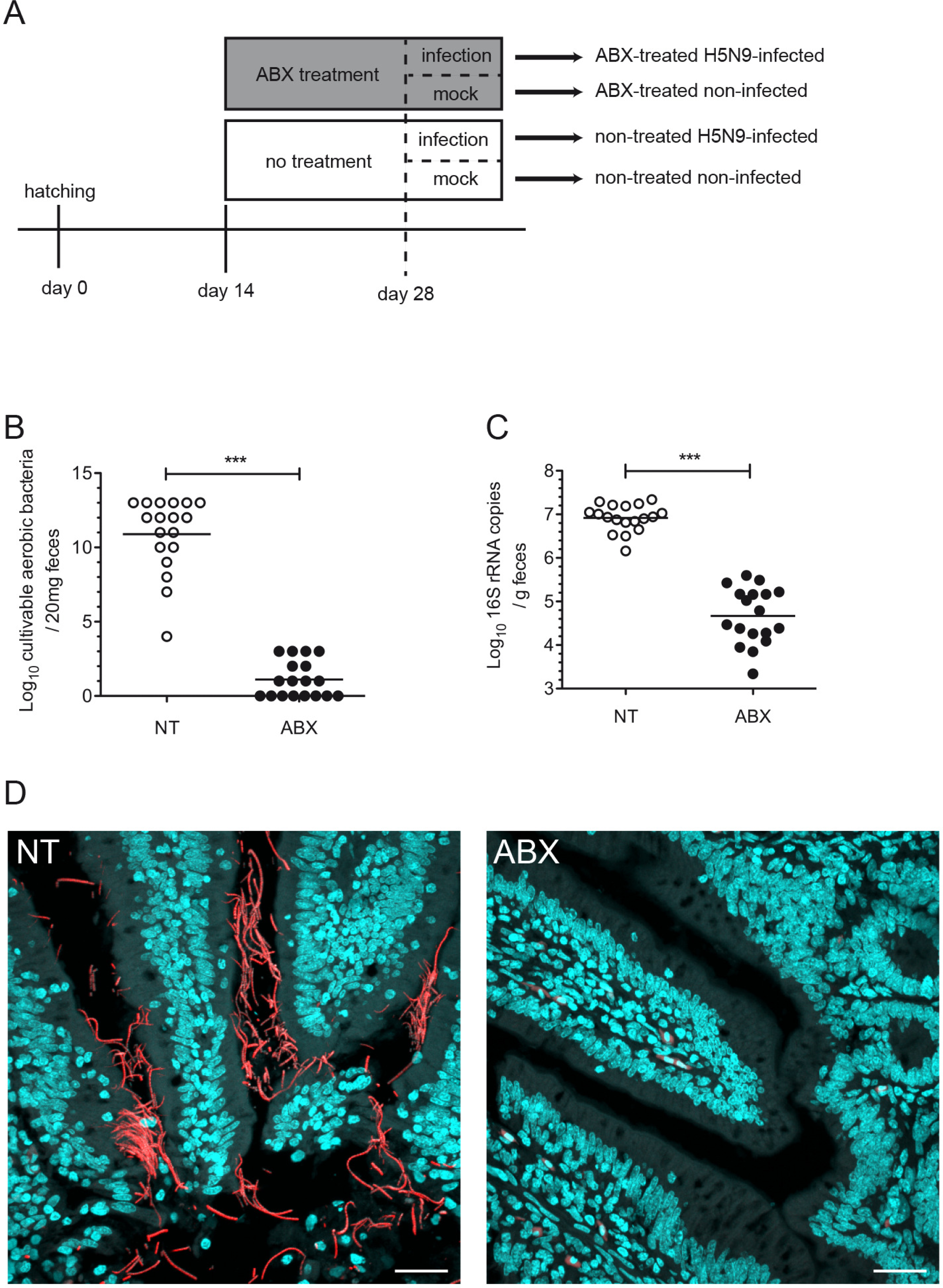
Antibiotic treatment for microbiota depletion. A) Treatment timelines for non-treated and antibiotics (ABX)-treated ducks. Ducks received the ABX cocktail in drinking water. Fecal bacterial loads were determined at day 28 after two weeks of ABX treatment and prior to H5N9 virus inoculation. B) Cultivable aerobic bacterial load per 20mg of feces. Feces harvested on day 28 from non-treated (NT) or ABX-treated ducks were diluted in bacterial broth and grown in aerobic liquid cultures under agitation. C) 16S rRNA gene copies per g of feces. DNA was extracted from feces harvested on day 28 and analyzed by qPCR using. D) Ileum sections from NT or ABX-treated ducks were subjected to fluorescent in situ hybridization (FISH) with eubacterial 16S-rRNA-specific Alexa594-labelled probe and stained with DAPI. Scale bar=25µM. ***p<0.001.

### Impact of intestinal microbiota depletion on influenza virus replication

We inoculated ducks via the ocular, nasal and tracheal route with 3.6×10^6^ egg infectious dose 50 (EID_50_) of the A/Guinea Fowl/France/129/2015(H5N9) HPAIV virus. ABX-treated ducks remained under the same ABX-treatment for the whole duration of the experiment. Ducks did not show any clinical signs upon H5N9 infection in the non-treated and ABX-treated group.

We evaluated the level of virus excretion by quantifying viral nucleic acids by RT-qPCR from tracheal and cloacal swabs. Tracheal shedding was equivalent between non-treated and ABX-treated ducks (Fig. 2A). By contrast, cloacal shedding was higher in ABX-treated animals at 3 and 5 days post infection (dpi), with a hundred-fold difference at 3 dpi (Fig. 2B).

**Figure 2.**
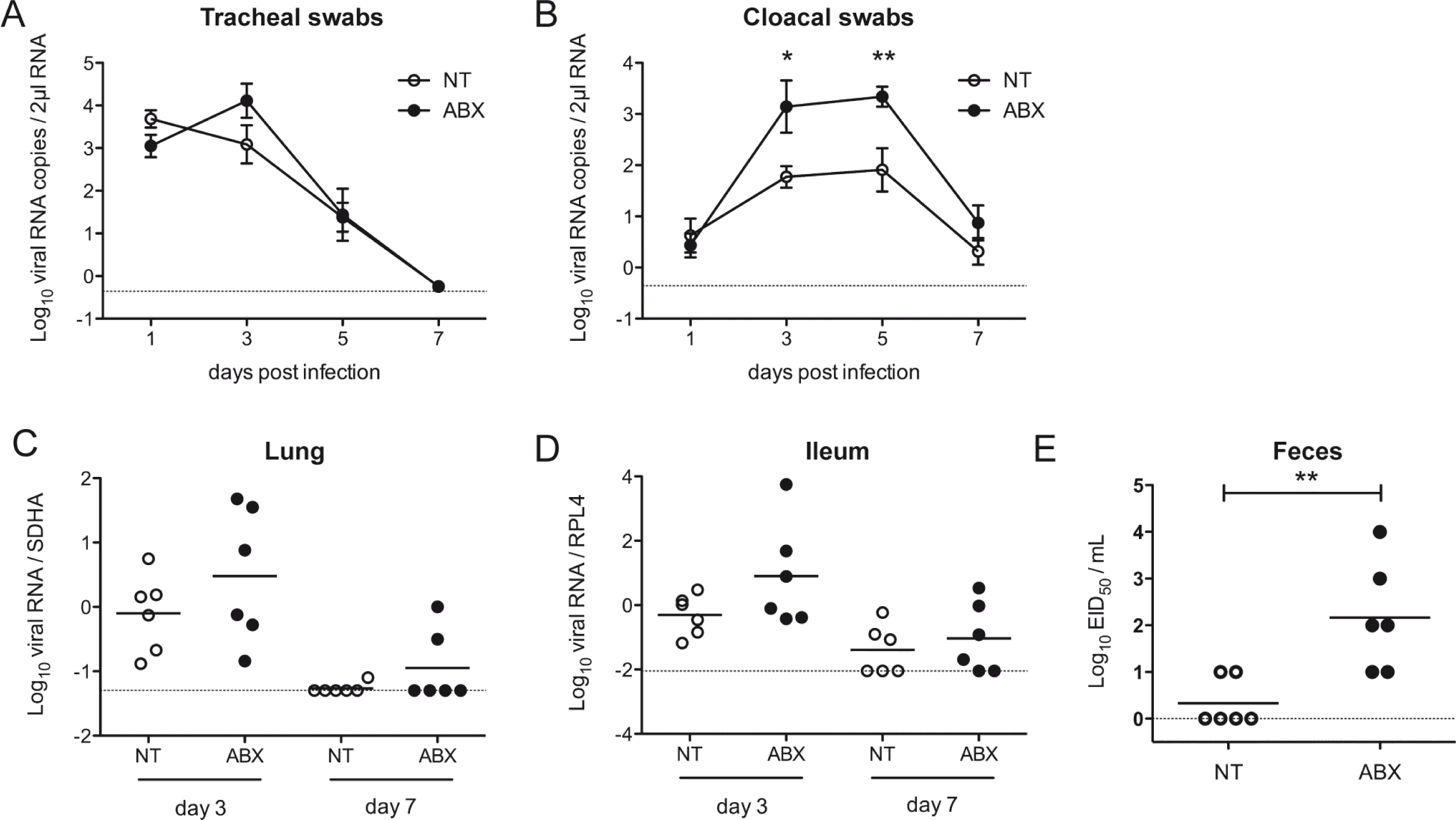
Consequences of microbiota depletion on H5N9 virus replication. NT or ABX-treated ducks were inoculated via the ocular, nasal and tracheal route with 3.6×10^6^ EID_50_ of the H5N9 HPAIV virus. Virus replication was analyzed by quantifying viral RNA by RT-qPCR from RNA extracted from tracheal swabs (A) or cloacal swabs (B). The curves represent the mean viral RNA load and error bars correspond to the standard error of the mean (SEM). Viral RNA load was analyzed from total RNA extracted from the lungs (C) and the ileum (D). Viral RNA was normalized with mRNA of the SDHA housekeeping gene in the lungs or the RPL4 housekeeping gene in the ileum. Each dot represents an individual value and the horizontal bar corresponds to the mean. (E) Viral load in the feces of animals autopsied at 3 dpi was titrated on chicken embryonated eggs and viral titers expressed as EID_50_. Each dot represents an individual value and the horizontal bar corresponds to the mean. The dotted line represents the limit of detection for each experiment. *p<0.05, **p<0.01

We then analyzed viral load in organs harvested from animals autopsied at 3 and 7dpi. Viral nucleic acid was detected in the brain and in the spleen of infected animals, as expected with a HPAIV, with no difference observed between ABX-treated-ducks and non-treated ducks (data not shown). In the lungs and ileum, viral nucleic acid load was increased in ABX-treated-ducks compared to non-treated ducks at 3 dpi (Fig. 2C&D), but the differences did not reach statistical significance. Finally, we measured infectious virus particles from fresh feces collected from individual ducks at 3 dpi. We detected a significantly higher number of infectious virus particles in feces from ABX-treated-ducks compared to non-treated ducks (Fig. 2E), thus confirming that ABX-induced depletion of the intestinal microbiota is associated with an increase of influenza virus fecal shedding in ducks.

### Impact of intestinal microbiota depletion on the respiratory and intestinal epithelia

Histopathological analysis performed at 3 and 7 dpi revealed a diffuse to multifocal subacute tracheitis in ducks infected with H5N9 virus, which was equivalent in ABX-treated and non-treated animals. Inflammatory cellular infiltrates in the lamina propria were variably composed of mononuclear cells (lymphocytes, plasmocytes and macrophages) and a few heterophils. We observed mild focal necrosis and exfoliation of the superficial mucosal epithelium, associated with loss of ciliature and regenerative epithelial hyperplasia. Histological scores were low for infected animals, yet different from non-infected animals (Fig. 3A). Immunohistochemically, viral antigen was detected in epithelial cells of the trachea, similarly in ABX-treated and non-treated ducks infected with H5N9 virus (Fig. 3B). In the ileum, we did not detect significant lesions associated neither with ABX-treatment nor H5N9 infection (data not shown). Viral antigen was detected in differentiated epithelial cells of the ileum, as well as in desquamated cells in the intestinal lumen (Fig. 3C).

**Figure 3.**
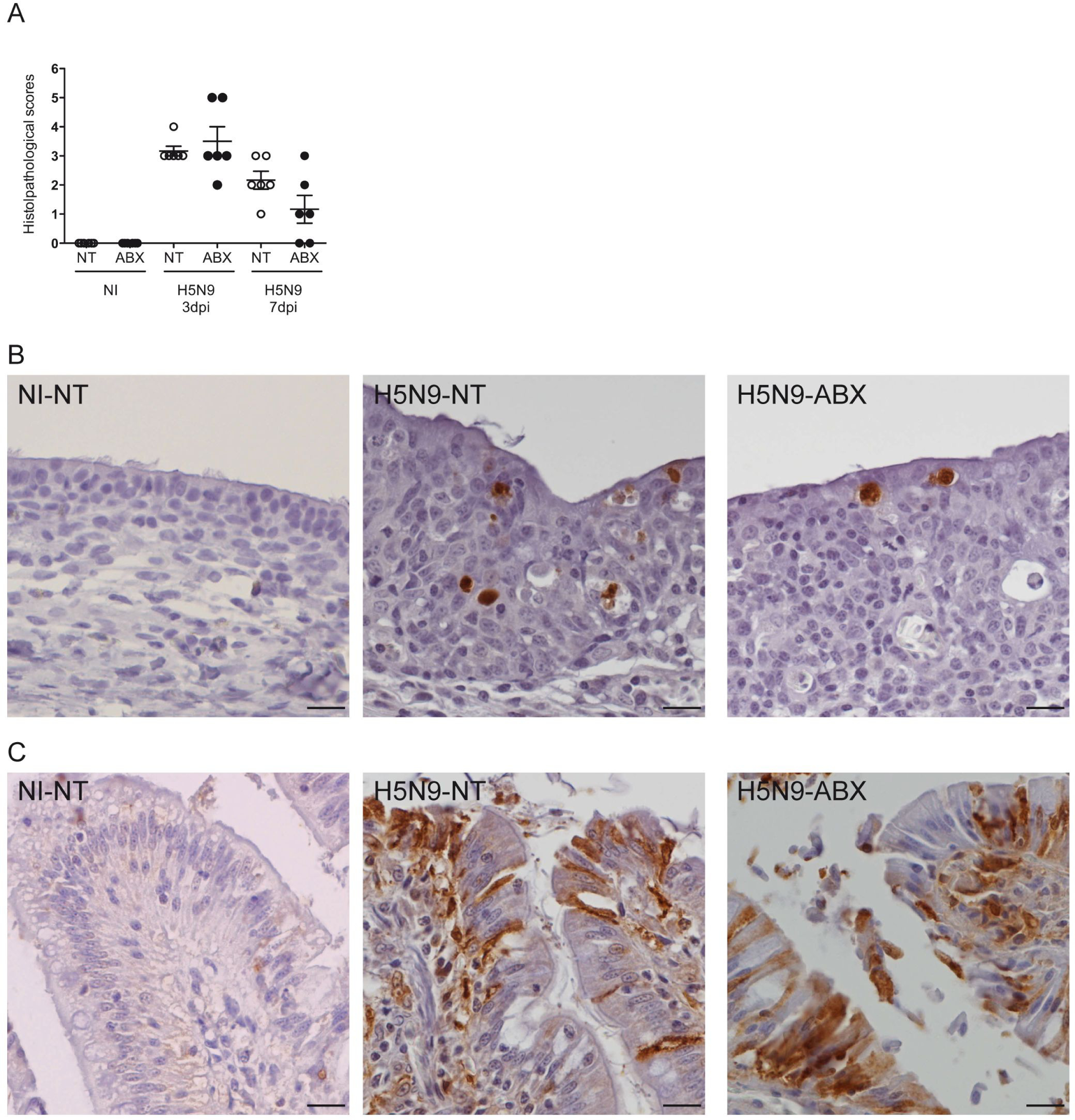
Histopathological analysis of H5N9 infection in NT and ABX-treated ducks. Histopathological analysis was performed on tracheal and ileum samples from non-infected (NI) or H5N9-infected ducks that were either non-treated (NT) or treated with ABX. (A) Histological scoring of haematoxylin and eosin stain colored tracheal sections. Each dot represents an individual value, the horizontal bar corresponds to the mean and error bars correspond to the SEM. Immunohistochemical anti-NP staining of haematoxylin counterstained tracheal (B) or ileum sections (C). Scale bar=10µm.

To gain further insight into the potential consequences of increased intestinal replication of H5N9 associated with depletion of the bacterial flora, we analyzed the expression of tight junction genes on ileal samples. We observed a significant increase in Claudin-1 mRNA levels in ducks infected with H5N9 at 3 dpi.

However, this increase of Claudin-1 mRNA expression was abrogated in H5N9-infected ducks treated with ABX (Fig. 4A). Similar results were obtained with Zonula occludens-1, another tight junction gene, for which a twofold non-statistically significant upregulation was observed specifically in non-treated H5N9 infected ducks at 3dpi (Fig. 4B). Thus, Claudin-1 and Zonula occludens-1 upregulation in the ileum in response to H5N9 infection may be dependent on the presence of a bacterial flora in the intestine. We also analyzed the expression of the gene encoding Mucin 2, a secreted high molecular weight glycoprotein, which is a component of the intestinal mucus. Mucin 2 expression was reduced specifically in H5N9-infected ducks treated with ABX (Fig. 4C). These results indicate that depletion of the bacterial flora may compromise intestinal integrity in response to HPAIV infection. To determine if depletion of the intestinal bacterial flora could affect influenza virus receptor expression, we generated fluorescent trimeric H5 HA to detect H5N9 virus receptor expression levels and distribution on tissue sections (20). The intensity and distribution of H5N9 receptors were similar in the trachea (Fig 5A) and intestine (Fig 5B) of ABX-treated and non-treated ducks. This result suggests that the increased replication of H5N9 in the intestinal tract of ABX-treated ducks is not due to a difference in virus receptor expression.

**Figure 4.**
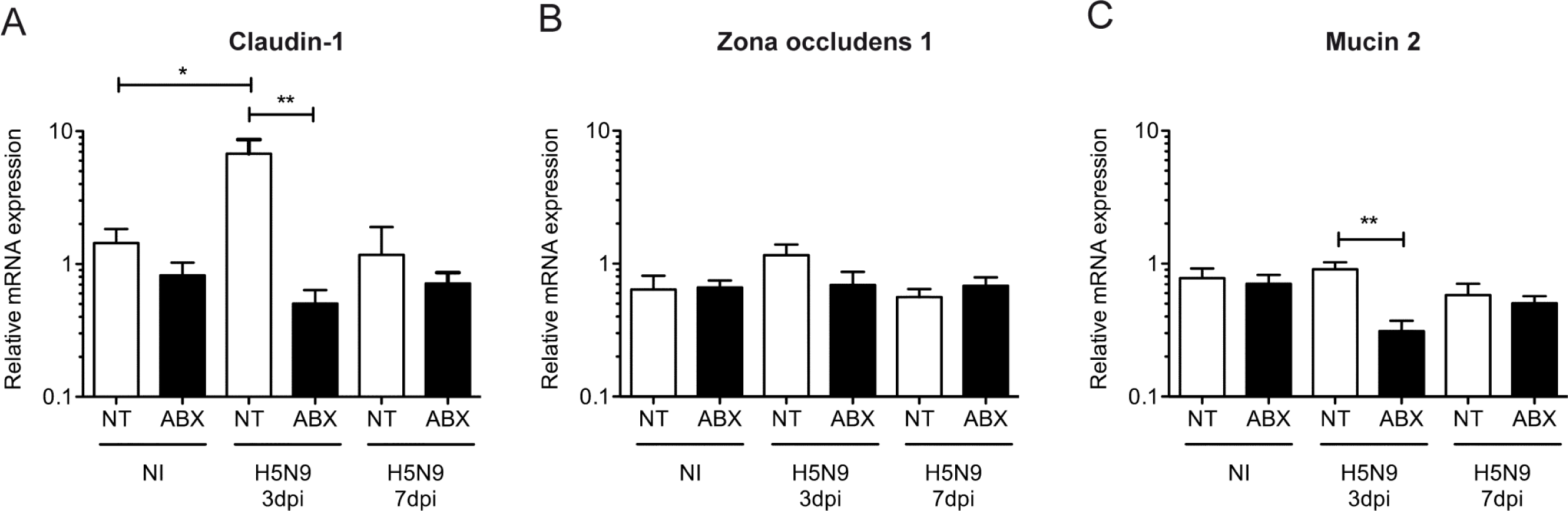
Analysis of intestinal integrity following H5N9 infection in NT and ABX-treated ducks. Analysis of mRNA gene expression from total RNA extracted from the ileum of non-infected (NI) or H5N9-infected ducks that were either non-treated (NT) or treated with ABX. RT-qPCR analysis of Claudin-1 (A), Zona occludens 1 (B) and Mucin 2 (C) mRNA levels in the ileum of ducks. mRNA levels were normalized to RPL4 levels. Results are expressed as means ± SEM. *p<0.05, **p<0.01

**Figure 5.**
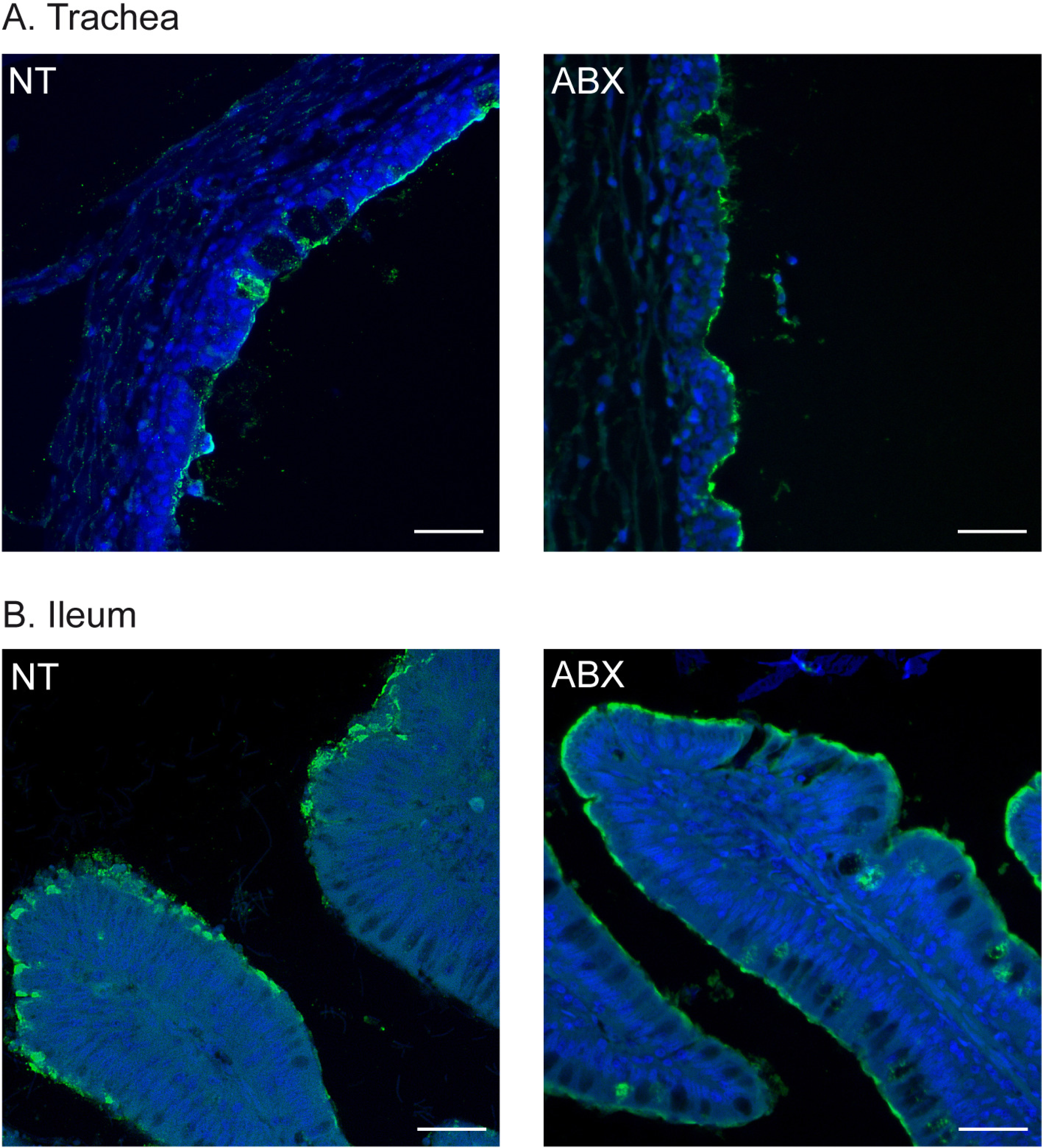
Analysis of H5N9 virus receptor expression. Staining of trachea (A) and ileum (B) from non-treated (NT) or ABX-treated ducks with trimeric HA from the H5N9 virus, precomplexed with α-strep-tag-488 conjugated mouse antibody and goat-α-mouse-488 conjugated antibody. Tissue sections were stained with DAPI (blue signal). Scale bar=25µm.

### Impact of intestinal microbiota depletion on the antiviral immune response

We then evaluated the impact of ABX-mediated intestinal microbiota depletion on the innate antiviral immune response. In the lungs, we observed an upregulation of IFN-α and IFN-β mRNA expression in H5N9-infected ducks at 3 dpi (Fig. 6A). This was associated with an upregulation of Interferon Induced Protein With Tetratricopeptide Repeats 5 (IFIT5), Oligoadenylate Synthetases-like (OAS-L) and Ubiquitin Specific Peptidase 18 (USP18) mRNA expression (Fig. 6A). IFIT5, OAS-L and USP18 are type I IFN-induced genes whose expression is a good indicator of the amount of type I IFN active locally. Interestingly, ABX-treatment was associated with a modest upregulation of IFN-α/β mRNA and type I IFN-induced genes expression regardless of the infection status.

**Figure 6.**
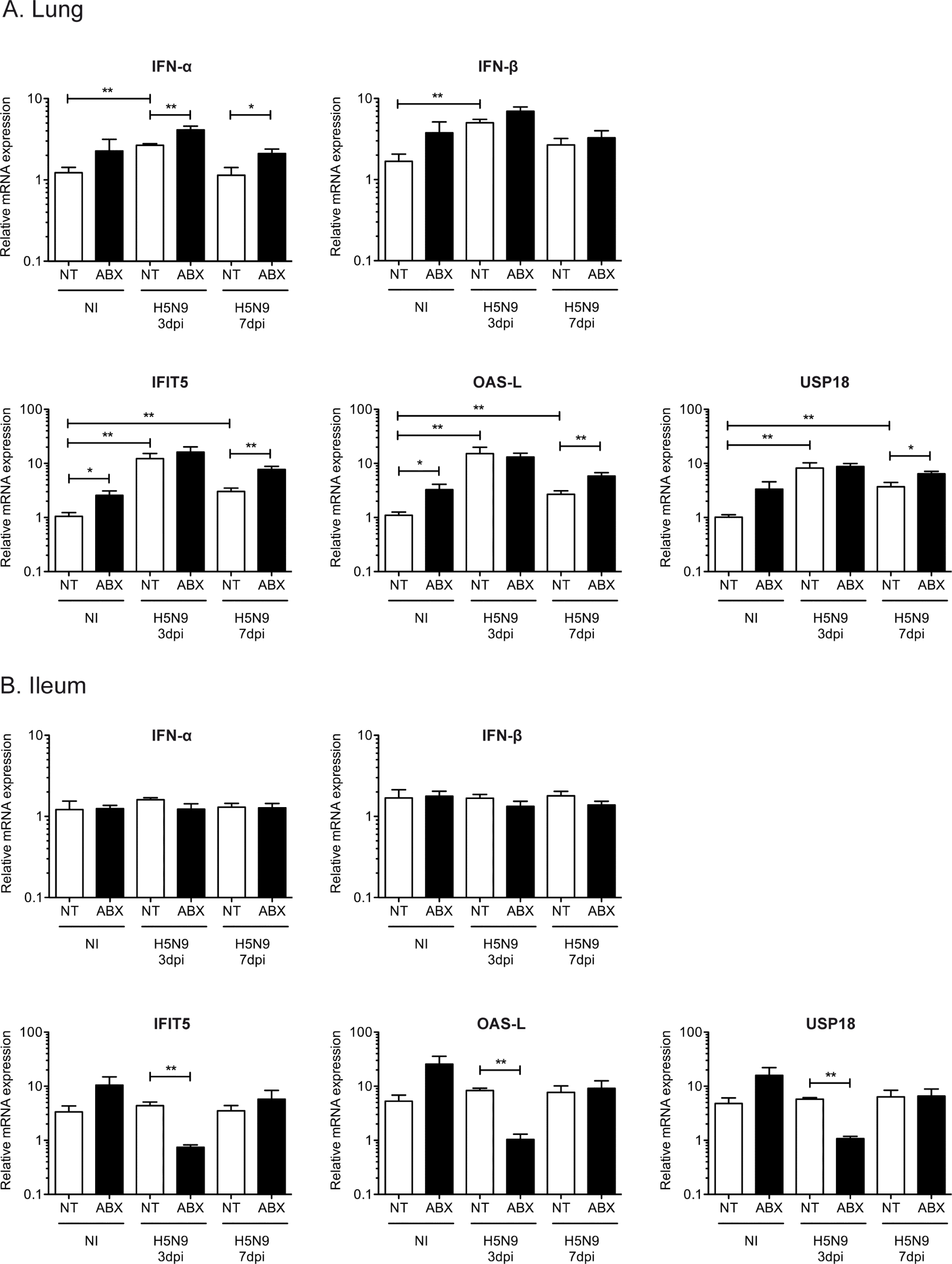
Analysis of the type I IFN immune response following H5N9 infection in NT and ABX-treated ducks. Analysis of mRNA gene expression from total RNA extracted from the lung (A) and ileum (B) of non-infected (NI) or H5N9-infected ducks that were either non-treated (NT) or treated with ABX. The mRNA expression levels of IFN-α, IFN-β and of three type I IFN-induced genes, IFIT5, OAS-L and USP18 were analyzed by RT-qPCR. mRNA levels were normalized to the geometric means of RPL4/SDHA mRNA levels in the lung and RPL4/RPL30 mRNA levels in the ileum. Results are expressed as means ± SEM. *p<0.05, **p<0.01

In the ileum, we did not detect significant upregulation of interferon-α (IFN-α) or IFN-β mRNA expression in ducks infected with HPAIV, regardless of the status of the intestinal microbiota (Fig. 6B). However, we observed an upregulation of type I IFN-induced genes in non-infected ABX-treated ducks (Fig. 6B), but the differences did not reach statistical significance. Interestingly, type I IFN-induced genes expression was significantly reduced at 3 dpi in H5N9-infected ABX-treated ducks compared to H5N9-infected non-treated ducks (Fig. 6B).

In order to evaluate if microbiota depletion could affect the antiviral humoral response in the intestine, we quantified the IgA expression levels in intestinal samples. We observed strongly reduced IgA heavy chain constant region mRNA expression in the intestine of ABX-treated ducks, regardless of the H5N9 infection status (Fig. 7A). We next analyzed the anti-H5 immunoglobulins levels in fecal samples that were available from non-infected ducks, as well as from H5N9-infected ducks at 3 dpi. We detected an increase in the anti-H5 immunoglobulins levels in feces collected from H5N9-infected non-treated ducks at 3 dpi, while no increase was observed in H5N9-infected ABX-treated ducks (Fig. 7B). Altogether these results indicate that increased H5N9-replication in the intestinal was associated with a reduced antiviral type I IFN-induced response, as well as with an impaired intestinal mucosal humoral immune response.

**Figure 7.**
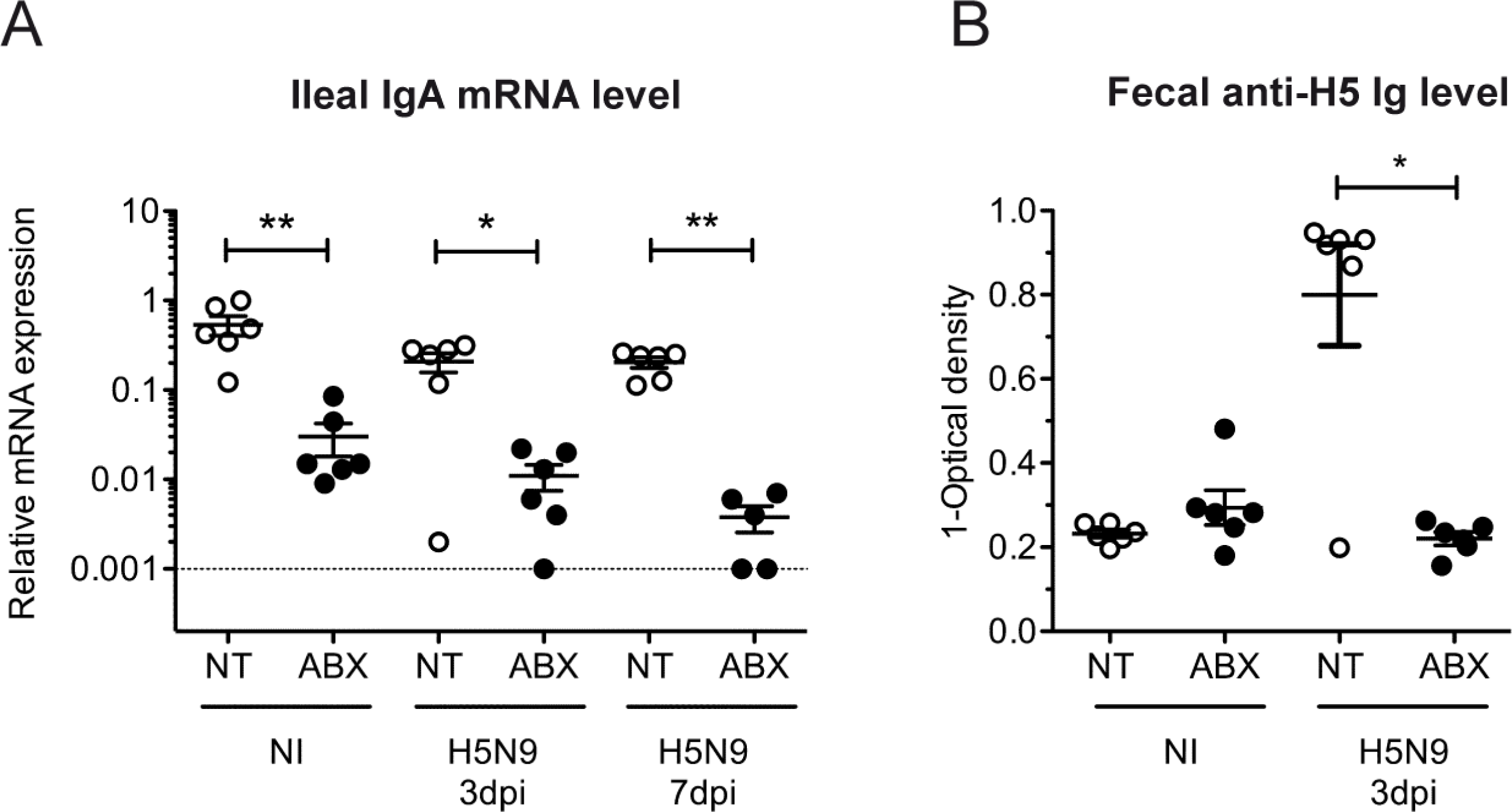
Analysis of the intestinal immunoglobulin production following H5N9 infection in NT and ABX-treated ducks. (A) IgA heavy chain constant region mRNA expression levels were analyzed by RT-qPCR from total RNA extracted from the ileum. IgA expression levels were normalized to RPL4 mRNA levels. (B) Fecal anti-H5 immunoglobulin levels were determined by a competitive anti-H5 ELISA performed on supernatants from fresh feces available from non-infected (NI) or H5N9-infected ducks at 3dpi. Each dot represents an individual value, the horizontal bar corresponds to the mean and error bars correspond to the SEM. *p<0.05, **p<0.01

## DISCUSSION

The A/Guinea Fowl/France/129/2015(H5N9) HPAIV virus studied here is a field isolate collected from a duck farm in France during the 2015-2019 H5Nx epizootics. The first HPAIV detected during this epizootic was of the H5N1 subtype. Genetic analyses revealed that the HA was independent from the Gs/Gd/1/96-like H5 lineage and thus represented a new HPAIV emergence event (21). The virus has undergone multiple reassortment events during its circulation in France leading to the co-circulation of H5N1, H5N2 and H5N9 HPAIV subtypes (22). Field observations provided evidence that clinical signs depended on the HPAIV subtype and on the host species. Gallinaceous poultry exhibited a moderate increase in mortality, while waterfowl mostly showed no clinical signs of infection (22). Here, we performed the first experimental H5N9 HPAIV infection in ducks (*Anas platyrhynchos*), which confirmed that H5N9 HPAIV infection in ducks was not associated with clinical signs and only produced mild lesions detected microscopically in the respiratory tract.

The contribution of the gut microbiota to the control of influenza virus replication has been studied in mice, as well as in chickens infected with a LPAIV (13, 14, 16–19, 23). How the gut microbiota modulates HPAIV infection, or more generally influenza virus infection in ducks has to our knowledge never been investigated. Here we provide evidence that ABX-mediated depletion of the intestinal microbiota caused an increase in H5N9 HPAIV replication in the digestive tract of ducks and was associated with a reduction of the antiviral immune response in the intestine. Similar results were observed in the digestive tract of chickens infected with a LPAIV (18, 19). Depletion of the gut microbiota in mice and chicken was associated with an increase in influenza virus replication in the respiratory tract (13, 14, 18, 19). By contrast, we did not detect a significant increase in influenza virus replication in the respiratory tract of ducks suggesting that this difference could be due to the viral strain used or host factors.

Upon depletion of the gut microbiota with ABX, we observed a reduction of the type I IFN-induced gene expression, as well as a reduction of the humoral response in the intestine following influenza virus infection. The gut microbiota has been shown to stimulate the immune response to pathogens most likely through the constant low-level exposure of epithelial cells and immune cells present in the intestinal mucosa to microbiota-derived pathogen-associated molecular patterns, as well as to microbial metabolites (23–26). This constant low-level stimulation is thought to maintain the immune system in an optimal state of reactivity allowing it to respond to pathogens via a timely innate and adaptive immune response (17, 27, 28). In particular, the microbiota is required for IgA production in the intestine (29, 30).

While we detected a significant upregulation of type I IFN-induced genes expression in the respiratory tract of ducks infected with the H5N9 HPAIV, no upregulation could be detected in the digestive tract following H5N9 HPAIV infection. As a significant reduction of type I IFN-induced genes expression was observed at 3 dpi in H5N9-infected ABX-treated ducks, the expression of type I IFN-induced genes in the digestive tract is not at its minimal level in non-infected ducks. These observations thus suggest that the basal expression level of type I IFN-induced genes in the digestive tract of ducks may be constitutively high and could contribute to the efficient control of influenza virus replication in ducks through early inhibition of the viral life cycle.

Interestingly, type I IFN-induced gene expression was increased in the digestive and respiratory tract of non-infected animals treated with ABX, compared to non-treated non-infected animals. The ABX cocktail used in our experiments contained gentamicin, a member of the aminoglycoside class of ABX, which have been shown recently to induce type I IFN-induced gene expression through a toll-like receptor 3 dependent pathway, independently of the microbiota (31). However, aminoglycosides stimulated type I IFN-induced gene expression locally and did not lead to increased type I IFN-induced gene expression in the respiratory tract when administered orally (31). Therefore, in our experiments, how the ABX-cocktail led to an increase of type I IFN-induced gene expression remains unknown.

In association with the respiratory tract and internal organs, the digestive tract represents a major site of HPAIV replication and excretion in birds (7, 8, 11). Moreover, the digestive tract is the main site of LPAIV replication in ducks (9, 32–34). Thus, avian influenza viruses can be considered as adapted to the digestive tract in ducks. A number of studies have shown that the gut microbiota promotes infection by enteric viruses such as poliovirus, norovirus or rotavirus, through a variety of processes including promotion of virus stability, attachment and entry, or evasion of the immune response (35–39). Our results thus highlight another type of interaction between an enteric virus and the microbiota as we observed impairment of avian influenza virus replication in the intestine of ducks harboring an intact microbiota compared to ABX-treated ducks.

Increased influenza virus replication and excretion in chickens (18, 19) and ducks treated with ABX raises questions about the risks associated with ABX treatments administered to farmed animals. However, the ABX treatment used to deplete the gut microbiota in our study and in the vast majority of experimental studies consists of a broad-spectrum ABX cocktail composed of six antimicrobial molecules, including molecules which are not allowed to be used in veterinary medicine as their use is restricted to human medicine in hospital settings or to research. ABX-treatments given to farmed animals mostly consist of one or two antimicrobial molecules and therefore should not have such a profound effect on the microbiota composition. Considering that ABX consumption for food animal production is forecasted to increase worldwide (40), further studies are needed to investigate if narrow spectrum ABX treatments given to farmed animals could also have consequences on virus replication and spread. In that respect, it is crucial to keep in mind that ABX-treatments administered to farmed animals are critical to the well-being of animals and to the economic profitability of the vast majority of farms worldwide (41–43). ABX treatments are not the only cause of microbiota dysbiosis in food animals. Nutrition and environmental parameters at birth and during rearing have also been shown to be associated with microbiota dysbiosis (44, 45). In addition, microbiota dysbiosis can also indirectly promote the spread and burden of infectious diseases by impairing the response to vaccines (46–48). In conclusion, further studies are needed to evaluate the consequences of microbiota dysbiosis and of possible interventions with probiotics on the burden of infectious diseases (47, 49).

## MATERIAL AND METHODS

### Ethics statement

This study was carried out in compliance with European animal welfare regulation. The protocols were approved by the Animal Care and Use Committee “Comité d’éthique en Science et Santé Animales – 115”, protocol number 13205-2018012311319709.

### Animals and antibiotic treatment

One day-old Pekin ducklings (*Anas platyrhynchos domesticus*, ST5 Heavy) were obtained from a commercial hatchery (ORVIA - Couvoir de la Seigneurtière, Vieillevigne, France). The study employed only female birds to minimize between-individual variability arising from sex. Birds were fed *ad libitum* with a starter diet. Ducks were housed for two weeks in a litter-covered floor pen in a Biosafety level II facility at the National Veterinary School of Toulouse, France. They were then transferred into a Biosafety level III facility, equipped with poultry isolators (I-Box, Noroit, Nantes, France) that were ventilated under negative pressure with HEPA-filtered air. A total of 36 ducks were randomly assigned to four groups: 6 non-treated non-infected animals (NT, NI), 6 ABX-treated non-infected animals (ABX, NI), 12 ABX-treated infected animals (ABX, H5N9) and 12 non-treated infected animals (NT, H5N9). Ducks of the ABX groups were treated through drinking water. ABX were administered at the following daily dosage: 80mg/kg vancomycin, 300mg/kg neomycin, 200mg/kg metronidazole, 0.2g/kg ampicillin and 24mg/kg colistin. To prevent fungal overgrowth, ABX-treated ducks also received 2mg/kg amphotericin-B daily. The composition and posology of the ABX cocktail are similar to the ones described in other publications (14, 18, 50, 51). Water was changed every two days. ABX treatment started 2 weeks prior to infection and continued for the whole duration of the experiment. No difference in water consumption was observed between non-treated and ABX-treated ducks.

### Virus and experimental infection

One day prior to infection, serum was collected from all the birds to ensure that they were serologically negative to influenza virus by using a commercial Influenza A NP Antibody Competition ELISA kit (ID Screen; ID-Vet, Montpellier, France) according to the manufacturer’s instructions. The A/Guinea Fowl/France/129/2015(H5N9) (Genbank MN400993 to MN401000) highly pathogenic avian influenza virus was propagated in 10-day-old SPF embryonated chicken eggs (INRA PFIE, Nouzilly, France). Infectious allantoic fluid was harvested 72h post-inoculation and titrated in 10-day-old SPF embryonated chicken eggs to determine the 50% egg infective dose (EID_50_)/ml using the Reed-Muench method. At 4 week-old, ducks were infected with 3.6×10^6^ EID_50_ of virus via the ocular, nasal and tracheal route. Ducks of the non-infected groups received the equivalent volume of allantoic fluid collected from non-infected embryonated chicken eggs. Birds were observed for 7 days and clinical signs were recorded. Tracheal and cloacal swabs were performed on all animals at days 0, 1, 3, 5 and 7 dpi. Half of the infected animals were euthanized and necropsied at 3 dpi and the other half at 7dpi.

### Analysis of bacterial depletion

Fecal samples were collected one day prior to infection by placing ducks in individual clean plastic crates. A fraction of the fecal samples was used directly for bacterial culture in order to determine the amount of cultivable aerobic bacteria in feces. For this purpose, 100mg of fecal samples were homogenized in 200µl PBS and centrifuged at 150g at 4°C for 2min to pellet large debris. Supernatants were transferred into new tubes and serially diluted in brain-heart infusion broth up to a 10^-16^ dilution. 50µl of each dilution was added into wells of 96-well plates containing 100µL brain-heart infusion broth. After 24h incubation at 37°C, the last positive well was used for calculation of the number of cultivable bacteria in feces. For 16S rRNA gene DNA quantification, microbial DNA was extracted from 500mg of stool (PureLink^TM^ Microbiome DNA Purifiation Kit, Invitrogen, Thermo Fisher Scientific Inc., Canada), according to the manufacturer’s instruction. qPCR was performed with 2µl of extracted DNA according to the manufacturer instructions (QuantiFast SYBR Green PCR, Qiagen, Canada) using 16S-V5-F (AGCRAACAGGATTAGATAC) and 16S-V5-R (TGTGCGGGCCCCCGTCAAT) primers to amplify the V5 region from 16S rDNA genes. Absolute quantification was performed using a standard curve based on a 10-fold serial dilution of plasmid containing the 16S-V5 region of E. coli. Fluorescence *in situ* hybridization (FISH) staining of the ileum was done as described previously (52). The tissue sections were incubated with the universal bacterial probe EUB338 (5’-GCTGCCTCCCGTAGGAGT-3’) (Eurogentec, Liège, Belgium) conjugated to Alexa Fluor 594. A ‘non-sense’ probe (5’-CGACGGAGGGCATCCTCA-3’) conjugated to Alexa Fluor 594, was used as a negative control. Tissue sections were mounted with Vectashield mounting medium containing DAPI. Images were acquired on a Zeiss LSM710 confocal microscope (Carl Zeiss MicroImaging GmbH, Jena, Germany) at the cellular imaging facility of the CPTP (Toulouse, France).

### Quantification of viral excretion

Cloacal and tracheal swabs were briefly vortexed in 500µl of sterile PBS and viral RNA were extracted from 140µl, according to the manufacturer’s instructions (QIAamp Viral RNA, Qiagen, Toronto, Canada). Influenza virus nucleic load was determined by qRT-PCR using influenza virus segment M specific primers (IAV-M; Table. 1) in 96-well plates according to the manufacturer instructions (OneStep RT-PCR, Qiagen, Toronto, Canada). Absolute quantification was performed using a standard curve based on a 10-fold serial dilution of a plasmid containing the A/Guinea Fowl/129/2015(H5N9) M gene. EID_50_ was determined by diluting 0.1g of feces into 100µl PBS containing antibiotics: vancomycin (500mg/l), neomycin (500mg/l), metronidazole (500mg/l), ampicillin (1g/l) and colistin (80mg/l) to avoid bacterial growth. We had first verified that the antibiotics did not inhibit viral replication in embryonated chicken eggs. Samples were centrifuged at 1000g at 4°C for 2min to pellet debris and supernatants were diluted in antibiotic-complemented PBS with a 10x dilution factor. For each sample, 100µl were injected in three 10-day-old SPF embryonated chicken eggs, which were incubated for 72h before allantoic fluids were harvested. Presence of virus was revealed by a hemagglutination test.

**Table.1:**
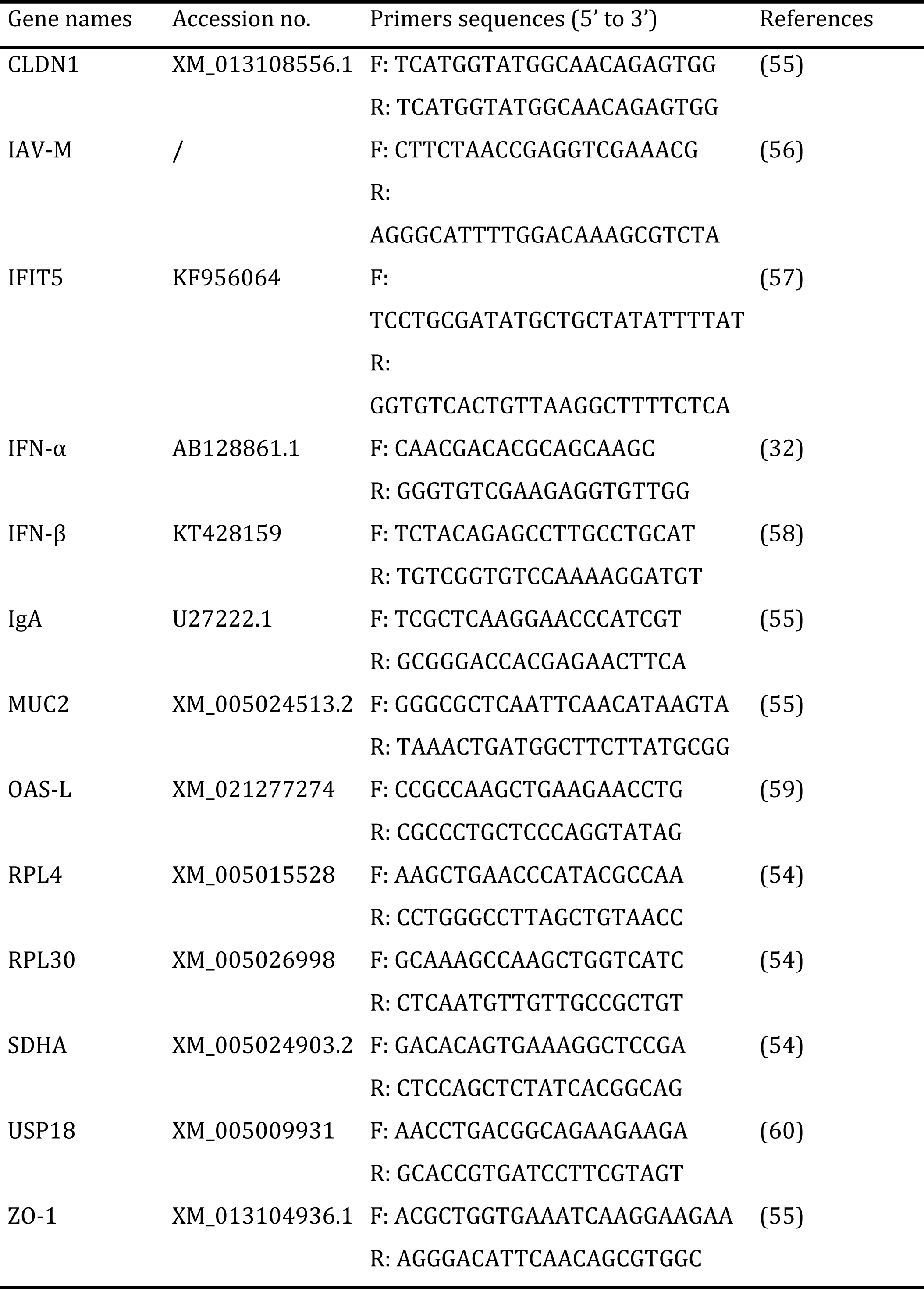
Primers used for qPCR.

### RNA extraction from tissue samples and complementary DNA synthesis

For each organ, 30 mg of tissue were placed in tubes with beads (Precellys lysis kit, Stretton Scientific Ltd, Stretton, UK) filled with 600µl of TRIzol reagent (Invitrogen, Carlsbad, CA, USA) and mixed for 30s at 6800 rpm three times in a bead beater (Precellys 24®, Bertin Technologies, Montigny-leBretonneux, France). Following TRIzol extraction, the aqueous phase was transferred to RNA extraction column and prcossed according to the manufacturer’s instructions (NucloeSpin RNA, Macherey-Nagel GmbH&Co, Germany). cDNA was synthesized by reverse transcription of 500ng of total RNA using both oligo(dT)18 (0.25µg) and random hexamer (0.1µg) and the RevertAid First Strand cDNA Synthesis Kit (Invitrogen, Thermo Fisher Scientific Inc., ON, Canada) according to manufacturer’s instruction.

### Quantitative PCR from tissue samples

Quantitative PCR for the analysis of host genes expression was performed in 384-well plates in a final volume of 5µl using a Bravo Automated Liquid Handling Platform (Agilent Technologies, Palo Alto, CA, USA) and a ViiA 7 Real-Time PCR System (Applied Biosystems, Foster City, CA, USA) at the GeT-TRiX platform (GénoToul, Génopole, Toulouse, France). Mixes were prepared according to the manufacturer instructions (QuantiFast SYBR Green PCR, Qiagen, Toronto, Canada) with 1µl of 1:20 diluted cDNA and a final concentration of 1µM of each primer (Table.1). Relative quantification was carried out using the 2^−ΔΔCT^ method (53) and the geometric means of two couples of housekeeping genes validated in specific duck tissues: RPL4/RPL30 and RPL4/SDHA for ileum and lung samples, respectively (54). Quantification of influenza virus nucleic acid load in tissues was performed in 96-well plates with 20µL final volume according to the manufacturer instructions (QuantiFast SYBR Green PCR, Qiagen, Toronto, Canada), 2µl of cDNA and a final concentration of 1µM of each primer. We used influenza virus segment M specific primers. For normalization, we used RPL30 for ileum samples and SDHA for lung samples, as they provided similar results to the geometric means of either RPL4/RPL30 or RPL4/SDHA. Reaction was performed on a LightCycler 96 (Roche, Mannheim, Germany) and relative quantification was carried out using the 2^−ΔΔCT^ method.

### Detection of anti-influenza H5 immunoglobulins in feces

For the detection of fecal anti-H5 specific immunoglobulins, we homogenized 100mg of feces in 150µl of sterile water and collected the supernatant after centrifugation at 150g at 4°C for 2min to pellet large debris. We then performed a commercial influenza H5 antibody competition ELISA (ID Screen; ID-VET, Montpellier, France). 50µL of fecal supernatant was used instead of diluted serum. The rest of the protocol was carried out according to the manufacturer’s instructions.

### Histopathological examination

All animals were subjected to a complete post-mortem examination. Tissue samples of trachea (one transversal section in the proximal portion and another one in the terminal portion), lungs, ileum, caecum, colon and brain were taken and stored in 10% neutral formalin. After fixation, tissues were processed in paraffin blocks, sectioned at 4μm and stained with hemalun and eosin (H&E) for microscopic examination. A board-certified veterinary pathologist who was blind to the experimental conditions assessed lesions histologically. Lesion intensity was graded as (0): no lesion, (1): minimal, (2): slight, (3): moderate, (4): marked or (5): severe.

### Immunohistochemistry

Immunohistochemistry was performed on paraffin-embedded sections of trachea with a monoclonal mouse anti-nucleoprotein Influenza A virus antibody (Argene, 11-030, pronase 0,05% retrieval solution, 10 minutes at 37°C: antibody dilution 1/50, incubation overnight, at 4°C). The immunohistochemical staining was revealed with a biotinylated polyclonal goat anti-mouse Immunoglobulins conjugated with horseradish peroxidase (HRP) (Dako, LSAB2 system-HRP, K0675) and the diaminobenzidine HRP chromagen (Thermo Scientific, TA-125-HDX). Negative controls comprised sections incubated either without specific primary antibody or with another monoclonal antibody of the same isotype (IgG2).

### H5 HA fluorescent tissue staining

A pCD5 plasmid was constructed with the HA sequence originating from A/guinea fowl/France/150207n/2015(H5N9) (Genbank KU320887.1), which has exactly the same amino acid sequence as the HA of A/Guinea Fowl/France/129/2015(H5N9). The plasmid encodes for a GCN4pI leucine zipper trimerization motif (**IL:**RMKQ**I**EDK**I**EE**IE**SK**QKKI**ENE**I**AR**I**KK, followed by a seven amino acid cleavage recognition sequence (ENLYFQG) of tobacco etch virus (TEV) and a sfGFP fused to a Strep-tag II (WSHPQFEKGGGSGGGSWSHPQFEK; IBA, Germany) C-terminally. The vector was transfected into HEK293S GNT1(-) cells (which are modified HEK293S cells lacking glucosaminyltransferase I activity (ATCC® CRL-3022™)) with polyethyleneimine I (PEI) in a 1:8 ratio (µg DNA:µg PEI). Tissues were stained with purified recombinant multimeric H5N9 as previously described (20).

### Statistical analysis

Statistical significance was determined by the Mann Withney test for individual time points. Statistical analyses were performed with Prism GraphPad software v5.01 (*p < 0,05; **p < 0,01; ***p < 0,001).

## ACKNOWLEDGEMENTS

We thank Jean-Luc Guérin and Luc Robertet (Université de Toulouse, ENVT, INRA, UMR 1225, Toulouse, France) for providing clinical samples that enabled the isolation of the A/Guinea Fowl/France/129/2015(H5N9) virus. We also thank Marie Souvestre and Luc Robertet for their assistance in the animal facility. We thank Sophie Allart and Astrid Canivet for technical assistance at the cellular imaging facility of INSERM UMR 1043, Toulouse. We thank the GeT-TRiX platform (GénoToul, Génopole, Toulouse, France) for access to the liquid handling platform and real-time PCR system.

This work was funded by the French National Agency for Research (ANR), project ANR-16-CE35-0005-01 Rule of Three to RV. PB is supported by a Ph.D. scholarship funded by the Region Occitanie (France) and by the Chaire de Biosécurité at the École Nationale Vétérinaire de Toulouse (French Ministry of Agriculture). RPdV is a recipient of an ERC starting grant (802780) and a Beijerinck Premium of the Royal Dutch Academy of Sciences (KNAW).

